# Ecological and metabolic signatures of biofilm viromes across a national river surveillance network

**DOI:** 10.64898/2026.10.26.761785

**Authors:** Susheel Bhanu Busi, Amy C. Thorpe, Jonathan Warren, Kerry Walsh, Daniel S. Read

**Affiliations:** UK Centre for Ecology & Hydrology, Wallingford, OX10 8BB, United Kingdom; Environment Agency, Horizon House, Bristol, BS1 5AH, United Kingdom

**Keywords:** virome, biofilm viruses, metagenomics, river surveillance, seasonality, river health

## Abstract

Benthic biofilms are biogeochemical hotspots in rivers, yet their endogenous viral communities remain poorly resolved at national scales. We analysed 447 biofilm metagenomes from 146 sites within England’s River Surveillance Network (RSN), sampled in spring and autumn from 2021 to 2023, integrating viral operational taxonomic units (vOTUs), water-quality gradients, predicted host associations and auxiliary metabolic genes (AMGs). We recovered 38,534 vOTUs, revealing extensive viral novelty, where 93% of the vOTUs could not be assigned to a known family. Viromes were dominated by Caudoviricetes and unclassified lineages and were extremely heterogeneous: no vOTU occurred in more than 8.3% of samples. Season, but not sampling year, structured virome composition (family-level PERMANOVA R^2^ = 0.050, p = 0.001; year p = 0.08), and within-site turnover between spring and autumn exceeded turnover between years. Bray–Curtis ordination linked virome structure weakly to pH, dissolved organic carbon, conductivity, nitrate-N and dissolved oxygen (envfit R^2^ ≤ 0.066), while threshold indicator taxa analysis (TITAN) identified vOTUs with abrupt change points along alkalinity, dissolved oxygen, nitrate-N and temperature gradients. Predicted host associations were concentrated in Pseudomonodota, Cyanobacteriota and Bacteroidota, with *Microcoleus*, *Chamaesiphon* and *Flavobacterium* among the dominant host genera. Virus–host networks were modular, and module abundances correlated with water quality variables (dissolved oxygen, pH, nitrate etc.), suggesting that environmental filtering acts partly through host ecology. Predicted virus–host associations formed a sparse network organised around a few highly connected viral lineages. Functional profiling showed that the biofilm virome was dominated by nucleotide metabolism, DNA methylation, phosphate-stress, glycan-modification and photosynthesis-related AMGs, with *nrdA*/*nrdB* ribonucleotide reductase as the dominant AMG signal. Few vOTUs (1.2%) were detected as proviruses, although provirus fractions were higher in selected host-linked lineages. Collectively, these results show that river biofilm viromes are seasonally structured ecological communities whose composition, host associations and metabolic potential reflect the physicochemical and biological state of rivers. We propose that biofilm viromes provide a complementary layer of information for understanding river health, microbial ecology and the propagation of anthropogenic pressures in freshwater systems.

## Introduction

River networks are active metabolic corridors that connect terrestrial landscapes, human infrastructure and downstream receiving waters.^1^ Within these systems, benthic biofilms form spatially structured microbial consortia on submerged surfaces and act as important sites of carbon transformation, nutrient cycling and contaminant processing. These biofilms comprise bacteria, archaea, algae, fungi and protists embedded within an extracellular polymeric matrix,^2^ creating a habitat in which microbial interactions are more persistent than in the water column.^3^ This is critical for river monitoring because biofilms do not simply record an instantaneous snapshot of the water column, but integrate environmental conditions,^4^ biological colonisation dynamics,^5,6^ and disturbance history^7^ over ecologically relevant timescales.^8^

While the bacterial component of rivers, including biofilms, has recently gained attention globally,^9,10^ the viral component of river biofilms remains comparatively understudied. Viruses influence microbial communities through host mortality, horizontal gene transfer, lysogeny and metabolic reprogramming during infection.^11^ In biofilms, these processes are likely to differ from planktonic environments because the matrix can restrict diffusion,^12^ increase local host densities and create steep microscale gradients in oxygen, nutrients and organic matter. These properties likely also favour distinct infection strategies and may allow viruses to influence both the stability and function of the biofilm. Additionally, viral auxiliary metabolic genes (AMGs), including genes involved in nucleotide metabolism, phosphate acquisition, carbohydrate transformation and photosynthetic maintenance,^13^ provide a mechanism through which viruses can alter host physiology^14^ and, in turn, biofilm biogeochemistry.

A central challenge in interpreting river viromes is distinguishing true ecological filtering from indirect associations with broad catchment descriptors. Wastewater treatment works, combined sewer overflows, agricultural land use and rainfall regimes can all affect river microbiomes, but their effects on viruses are likely to be mediated through more proximate drivers. These include but are not limited to nutrient concentrations, dissolved organic carbon, pH, oxygen availability, hydrological disturbance, chemical mixtures and the composition of bacterial, algal and eukaryotic host communities. This issue is particularly important for biofilm viruses because viral community composition is expected to reflect the ecology of available hosts as well as the direct introduction or removal of viral particles.

In this study, we characterise the ecology of river biofilm viromes across England, using samples collected through the national River Surveillance Network (RSN).^10^ We frame the study around two linked hypotheses. First, we hypothesise that river biofilms harbour a large and taxonomically under-characterised virome whose composition varies across time and season, and is associated with measured water quality gradients, reflecting both dispersal processes and local environmental context. Second, we propose that environmental filtering propagates through virus–host associations and viral functional traits, resulting in modular host–virus networks and non-random distributions of AMGs linked to river biofilm metabolism. By integrating vOTU composition, water chemistry, indicator responses, host prediction, network structure and AMGs, we move beyond a descriptive catalogue of river viruses towards a biological framework for understanding how viruses contribute to river biofilm ecology and river health.

## Materials and Methods

### Sampling and catchment metadata harmonisation

Benthic biofilm samples were collected by Environment Agency staff from RSN sites across England in spring (March–May) and autumn (September–November) from 2021 to 2023. At each site, biofilm was scraped with a sterile toothbrush from the upper surfaces of five stones taken from the riverbed. It was preserved in the field in 5 mL of nucleic acid preservation buffer (3.5 M ammonium sulphate, 17 mM sodium citrate, 13 mM EDTA)¹⁵, then transported to the EA National Laboratory (Starcross, Exeter), where samples were concentrated by centrifugation and frozen. DNA was extracted from 100 µL of each sample with the Quick-DNA Fecal/Soil Microbe kit (Zymo Research), modified as described in Thorpe *et al*.¹⁰. The modifications were DNA/RNA Shield as lysis buffer, bead-beating on a TissueLyser II (Qiagen) and proteinase K digestion at 65 °C for 20 min. DNA purity was checked on a NanoDrop 8000 and concentration was measured with the QuantiFluor ONE dsDNA kit on a BioTek Cytation 5. Libraries were prepared and sequenced by Novogene UK as 2 × 150 bp paired-end reads on an Illumina NovaSeq 6000 (S4 flow cell), targeting ≥ 10 Gb of raw data per sample.

In total, 450 biofilm metagenomes were sequenced with concomitant metadata collected including physicochemical data. Three metagenomes (L30_2021, L65_2021 and O12_2022) yielded no vOTU profile and were excluded. All remaining metagenomes were matched to the existing RSN sampling records. The final dataset comprised 447 metagenomes from 146 RSN sites (216 spring, 231 autumn; 172 in 2021, 177 in 2022 and 98 in 2023), and the same 447 samples were used in all analyses unless stated otherwise.

We focused our analyses on measured water quality variables available across the surveillance network, including orthophosphate-P, nitrate-N, ammoniacal-N, dissolved organic carbon (DOC), conductivity, pH, dissolved oxygen, alkalinity and water temperature (Supp. Table 1). These variables provide proximate ecological gradients likely to affect microbial hosts and, indirectly or directly, the biofilm virome. Seasonal differences in each variable were tested with Wilcoxon rank-sum tests with Benjamini–Hochberg (BH) correction.^16^

### Metagenome processing

Read processing, assembly and binning followed Thorpe et al.¹⁰ where adapters were trimmed and reads were quality-filtered (minimum Phred score 30) with Trim Galore v0.6.5. Additinally, reads mapping to the human reference genome (GRCh38) were removed. Each sample was assembled separately with MEGAHIT v1.2.9 and quality-filtered reads were mapped back to their own assembly with bbmap to produce per-sample BAM files. Metagenome-assembled genomes (MAGs) were recovered from the same assemblies with MetaBAT v2.15, MetaBinner v1.4.3 and CONCOCT v1.1.0. MAGs were dereplicated with dRep v3.2.2 at ≥ 95% average nucleotide identity (ANI) over ≥ 65% aligned fraction and quality-checked with CheckM2 v1.2.2 followed by taxonomic assignments using GTDB-Tk v2.3.2. This produced 820 medium-quality (≥ 70% completeness, < 10% contamination) and high-quality (≥ 90% completeness, < 5% contamination) species-level MAGs¹⁰.

### Virome bioinformatics pipeline

Viral sequences were identified and characterised from the per-sample assemblies and BAM files with a Snakemake workflow adapted from DiscoVir¹⁷, in which each step ran in its own conda environment (github.com/UKCEH-MolecularEcology/virome_pipeline; commit 941a718, archived on Zenodo; *see Code availability*). Viral contigs were identified with geNomad v1.13.0¹⁸ in end-to-end mode, with default post-classification filtering and score calibration (--enable-score-calibration). geNomad combines alignment-free neural-network and marker-gene classification to detect viral sequences and assign ICTV-based taxonomy. CheckV v1.1.1¹⁹ (database v1.5) estimated genome completeness and contamination and trimmed flanking host regions from proviruses. CheckV quality filter was not applied before clustering, so genome fragments were retained, but contigs shorter than 1 kb were discarded with seqtk v1.5.

Viral contigs from all samples were pooled and exact duplicates were removed with BBTools dedupe (BBMap v40.01). The remaining sequences were clustered with MMseqs2 v 17-b804f (--min-seq-id 0.95 -c 0.85 --cov-mode 1 --cluster-mode 2), so that each member shared ≥ 95% nucleotide identity with its cluster representative over ≥ 85% of its length. This approximates species-level viral operational taxonomic units (vOTUs). For vOTU abundance, reads in each sample’s BAM file were counted against geNomad-predicted coding sequences with VERSE v1.0. Counts were normalised to gene length (reads per kb) and scaled to one million per sample (hereafter CPM). Additionally, geNomad virus scores were used to assign confidence classes for visualisation (moderate < 0.90, high 0.90–0.97, very high 0.97–0.995, near-certain ≥ 0.995). vOTUs that geNomad identified as proviruses (integrated in a host contig) were classified as temperate. All other vOTUs are referred to as ‘not detected as provirus’, because fragmented genomes cannot be reliably assigned a lytic lifestyle.

For functional annotation, viral contigs (≥ 1 kb) were first processed with VirSorter2 v2.2.4 (--include-groups dsDNAphage,ssDNA,NCLDV, --min-score 0.5, --provirus-off, --prep-for-dramv). They were then annotated with DRAM-v v1.5²⁰ focusing on auxiliary metabolic genes (AMGs) involved in carbon, nitrogen, phosphorus, nucleotide, cell-envelope, redox and photosynthesis-related processes. Gene coordinates were transferred from the VirSorter2-trimmed contigs back to the original contigs with Liftoff v1.6.3, and AMG abundances were estimated with VERSE. For the AMG analyses we used the 401 samples with at least one DRAM-v-annotated AMG. Furthermore, hosts of prokaryotic viruses were predicted with iPHoP v1.4.²¹ iPHoP combines host-based signals (BLAST matches to host genomes, CRISPR-spacer matches, and WIsH, VirHostMatcher and PHP k-mer scores) with phage-based signals (RaFAH and similarity to reference phages of known host). Machine-learning classifiers integrate these into a confidence score for each virus–host genus pair. We used the iPHoP June 2025 database (Jun_2025_pub_rw), supplemented with the medium-quality or higher RSN MAGs from Thorpe *et al.*¹⁰ and retained predictions with a confidence score ≥ 90. vOTUs assigned to lineages that infect eukaryotes (1,716 vOTUs: the phyla Nucleocytoviricota, Peploviricota, Artverviricota, Cossaviricota, Pisuviricota and Lenarviricota, and the families Circoviridae, Nanoviridae, Lavidaviridae, Adintoviridae and Adenoviridae) were flagged and excluded from the host-association, network and lifestyle analyses. Smacoviridae, whose hosts are uncertain, were retained.

### Ecological and statistical analysis

All downstream statistical analyses were performed in R v4.3.3 using tidyverse, vegan v2.6-4,^22^ TITAN2 v2.4,^23^ igraph, ggraph, ggalluvial, patchwork and ggsci. vOTU abundances (counts per million, CPM) were converted to relative abundance and Hellinger-transformed. Community structure was assessed using Bray–Curtis dissimilarities and principal coordinates analysis (PCoA, Lingoes correction). Because vOTU-level communities were extremely sparse (91% of sample pairs shared no vOTU), analyses were repeated on communities aggregated to the lowest assigned viral taxon (family or higher; ‘family level’). Effects of season and year were tested by PERMANOVA (adonis2, 999 permutations)^24^ using marginal tests and a sequential season × year model, with a robustness check in which permutations were restricted within river catchments where homogeneity of dispersion was tested with betadisper. To quantify seasonal turnover directly, we compared Bray–Curtis dissimilarity between pairs of samples from the same RSN site collected in different seasons of the same year (seasonal; 180 pairs, 126 sites) with pairs from the same site and season in different years (interannual; 207 pairs, 67 sites) using Wilcoxon tests. Seasonally differential viral taxa (present in ≥ 10% of samples) and vOTUs (present in ≥ 2% of samples) were identified using Wilcoxon rank-sum tests with BH correction, and vOTU richness and Shannon diversity were compared between seasons and years. Environmental alignment with virome composition was evaluated using envfit (999 permutations; 439 samples with complete chemistry), with the four strongest water quality vectors plotted on the ordination.

Indicator responses were evaluated in two complementary ways. First, Spearman correlations between orthophosphate-P, the nutrient that differed the most between seasons and the relative abundance of each vOTU detected in ≥ 1% of samples (n = 242) were calculated. Second, threshold indicator taxa analysis (TITAN2) was used to identify vOTUs showing abrupt changes (change points) in abundance and frequency of occurrence along selected water quality gradients. Because no vOTU occurred in more than 8.3% of samples, TITAN2 was run on the 50 vOTUs detected in ≥ 2% of samples, using relative abundances as community input and including all samples with measurements for each environmental gradient, in line with our previous findings, the four gradients were selected a priori from Thorpe et al.^4^ (alkalinity, n = 402; dissolved oxygen, n = 441; nitrate-N, n = 440; water temperature, n = 441). Because TITAN gradients overlap with season, we report the overlap between indicator vOTUs and seasonally differential vOTUs. The analysis used 999 permutations and 999 bootstrap replicates, and vOTUs were retained as indicators when purity and reliability were both ≥ 0.95. These analyses were designed to identify environmentally responsive viral populations without assuming that catchment-pressure categories alone define the relevant ecological gradients.

Host–virus associations were characterised by linking predicted host phyla and genera to viral families and by constructing a bipartite network in which host genera and viral taxa were connected by edges weighted by the summed mean CPM of linking vOTUs. Network statistics were calculated using igraph, and Louvain modules^25^ were identified to evaluate compartmentalisation of host–virus associations; modules were numbered by size. Module-level viral abundances (summed CPM of vOTUs belonging to the viral taxa in each module) were correlated with water quality variables using Spearman correlations with BH correction to test whether network compartments were associated with environmental gradients. Lifestyle predictions were used to compare lytic and temperate vOTUs across genome-size distributions, major viral taxa and dominant predicted host genera. AMG profiles were summarised across the samples with DRAM-v AMG output, grouped into functional categories (nucleotide metabolism, DNA methylation/defence, phosphate stress, glycan/cell envelope, photosynthesis, lipid metabolism, redox/cofactor) and further resolved by encoding viral taxon to identify which metabolic functions dominated the biofilm virome.

## Results

### National-scale biofilm viromes reveal environmental gradients underpinning river microbial ecology

To establish the spatial and environmental context of the RSN virome dataset, we mapped 447 metagenome samples from 146 sites across England (see Methods for the three samples excluded). The dataset captured broad national coverage and included repeated spring and autumn sampling across 2021, 2022 and 2023 (Fig. 1a). This sampling design enabled us to test whether river biofilm viromes exhibit structured variation across seasons and environmental contexts, rather than representing a homogenised pool of viral particles transported through connected river systems.

**Figure 1.**
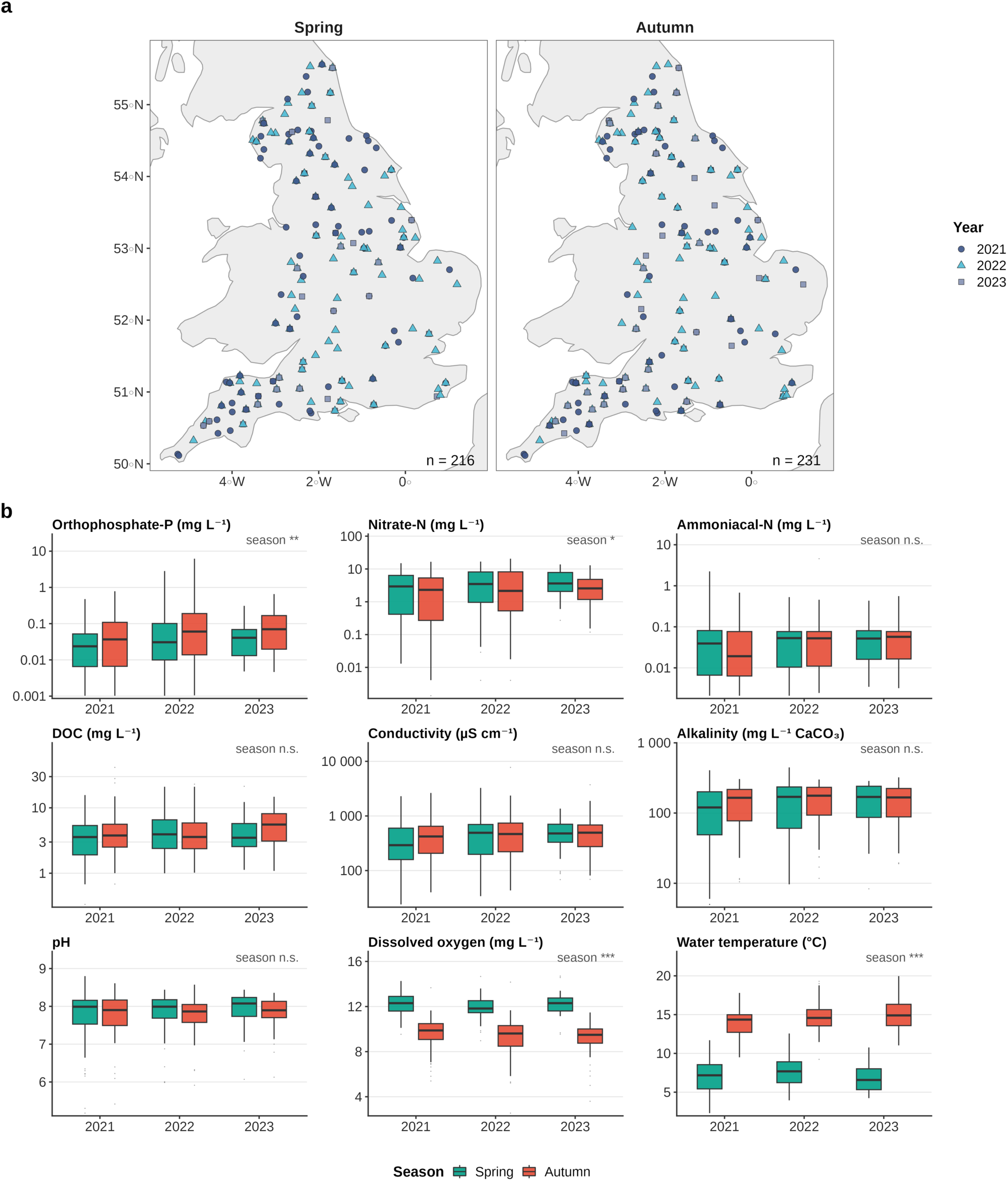
National sampling framework and water-quality context for the RSN virome. (a) Locations of the 447 biofilm metagenomes, shown separately for spring (n = 216) and autumn (n = 231) sampling, with sampling year indicated by colour and symbol. (b) Distributions of water-quality variables by year and season. Boxes show the median and interquartile range; whiskers extend to 1.5 × IQR. Skewed variables are shown on a log_10_ scale. Labels give the result of a Wilcoxon test between seasons (BH-adjusted; * q < 0.05, ** q < 0.01, *** q < 0.001; n.s., not significant).

Measured water quality data captured substantial physicochemical variation across the sampled sites, spanning several orders of magnitude for nutrients, DOC and conductivity (Fig. 1b and Supp. Table 1). Seasonal contrasts were pronounced for water temperature (median 7.3 °C in spring vs 14.5 °C in autumn) and dissolved oxygen (12.1 vs 9.6 mg L^−1^; both BH-adjusted q < 0.001), and significant but smaller for orthophosphate-P (higher in autumn; q = 0.003) and nitrate-N (higher in spring; q = 0.02), whereas ammoniacal-N, DOC, conductivity, alkalinity and pH did not differ between seasons (Fig. 1b). Because these variables influence microbial growth, biofilm metabolism and host availability, they provide more biologically interpretable axes for virome analysis than broad catchment pressure classes alone.

### River biofilms contain a highly novel and seasonally structured virome

Across the RSN biofilms, we recovered a large and taxonomically under-characterised viral community comprising 38,534 vOTUs, all of which were detected in the 447 analysed samples. Virome composition was consistently dominated by Caudoviricetes and unclassified viral lineages, with additional contributions from Adintoviridae, Mimiviridae, Smacoviridae, Autographiviridae, Phycodnaviridae, Crassvirales and other viral taxa (Fig. 2a). The virome was extremely heterogeneous among samples: most vOTUs were detected in a single sample, and no vOTU occurred in more than 8.3% of samples (Supp. Fig. S1c), and 91% of sample pairs shared no vOTU. Consequently, vOTU-level Bray–Curtis dissimilarities were close to saturation and season, although significant, explained only 0.3% of vOTU-level variation (PERMANOVA R^2^ = 0.003, p = 0.001), with no effect of year (p = 0.07) or season × year interaction (p = 1.0). When vOTUs were aggregated to the lowest assigned viral taxon (family or higher), season explained 5.0% of compositional variation (R^2^ = 0.050, p = 0.001), whereas year remained non-significant (R^2^ = 0.007, p = 0.08; interaction p = 0.79). Results were unchanged when permutations were restricted within rivers (season p = 0.001). At the vOTU level, mean distances to the seasonal centroids were virtually identical (0.704 in both seasons; betadisper p = 0.005), the significance reflecting the near-constant, saturated dissimilarities rather than a meaningful difference in spread. At the family level, spring viromes were more dispersed than autumn viromes (mean distance to centroid 0.25 vs 0.22; p = 0.001). Because the design was nearly balanced (216 spring vs 231 autumn samples), PERMANOVA is relatively robust to this heterogeneity,^26^ but the family-level seasonal effect is likely to reflect differences in both composition and dispersion.

**Figure 2.**
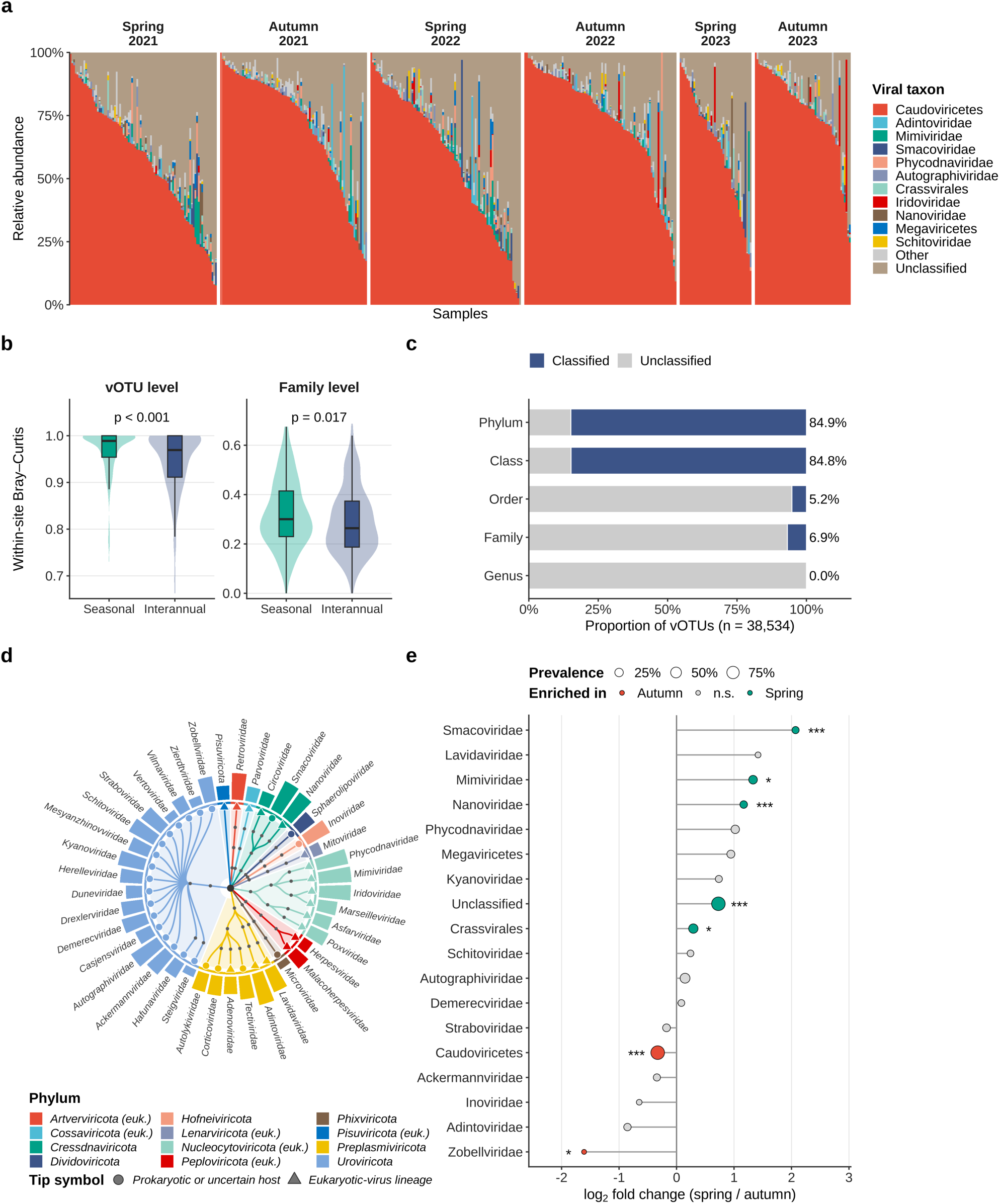
Composition, seasonal structure and taxonomic novelty of the RSN virome. (a) Relative abundance of the dominant viral taxa (family or lowest assigned rank) in each of the 447 samples, grouped by season and year and ordered by Caudoviricetes abundance. (b) Within-site Bray–Curtis dissimilarity between samples from the same RSN site collected in different seasons of the same year (seasonal) or in the same season of different years (interannual), at vOTU and family level; p values from Wilcoxon tests. (c) Proportion of the 38,534 vOTUs classified at each taxonomic rank. (d) Taxonomy-based circular dendrogram of classified viral lineages; tips are viral families (or the deepest classified rank), shaded sectors and the inner ring indicate phylum, and outer bars show log_10_(mean CPM + 1). Triangular tips and “(euk.)” in the legend mark lineages of viruses that infect eukaryotes. (e) Log_2_ fold change in mean relative abundance (spring/autumn) of viral taxa present in ≥ 10% of samples; point size shows prevalence and colour the season of enrichment (BH-adjusted Wilcoxon * q < 0.05, *** q < 0.001).

**Supplementary Figure 1.**
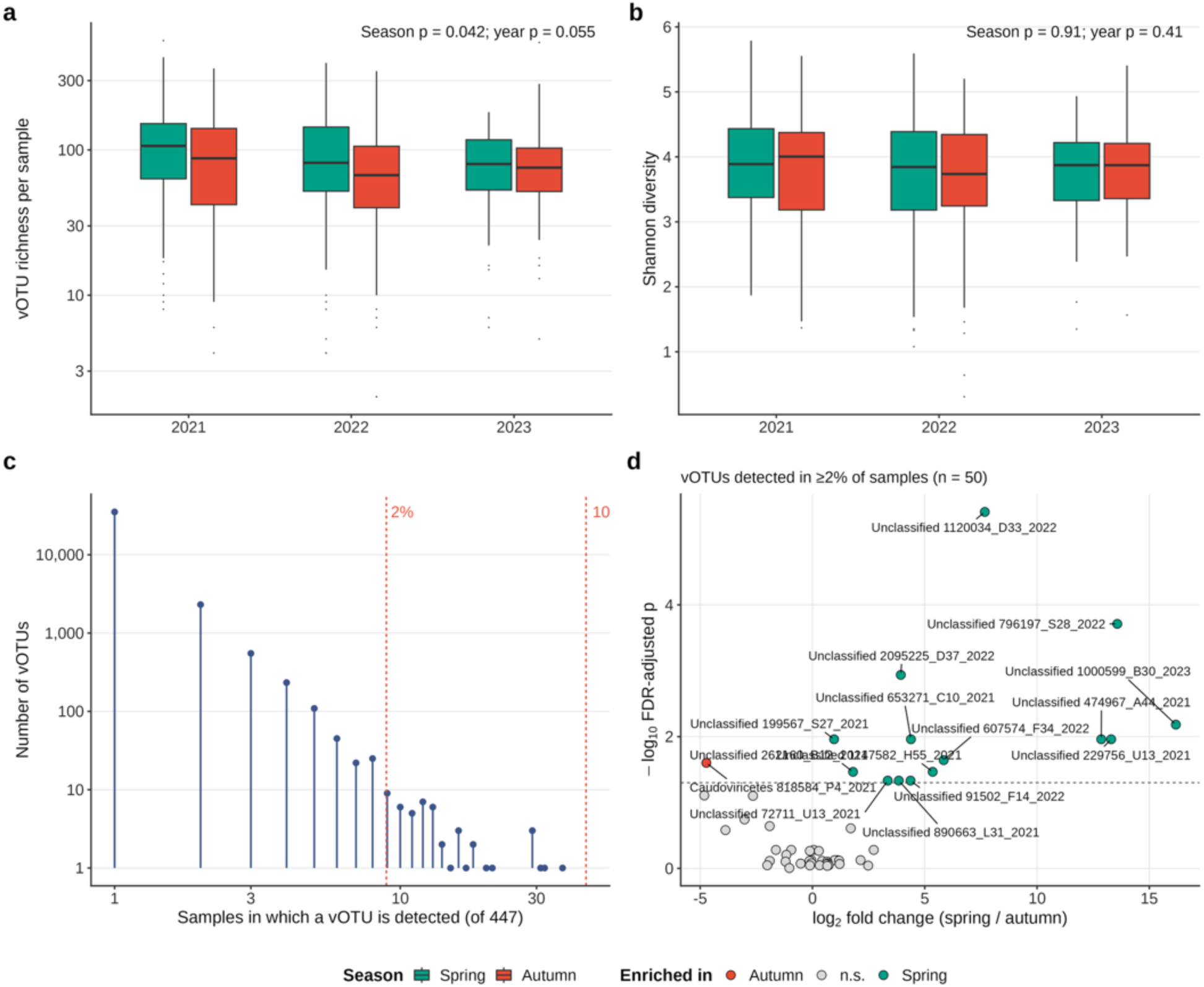
Virome metrics across the RSN. vOTU richness (a) and Shannon diversity (b) across the seasons and years. (c) Prevalence of vOTUs across 447 samples. Red dotted lines represent the 2% and 10% prevalence thresholds. (d) Differential abundance of vOTUs when comparing Spring to Autumn seasons, found in at least 2% of the samples. Points indicate vOTUs with an FDR-corrected significance of p < 0.05.

Seasonal turnover was also evident within sites. For RSN sites sampled more than once, vOTU composition differed more between spring and autumn of the same year than between the same season in different years (median Bray–Curtis 0.989 vs 0.969; Wilcoxon p < 0.001; Fig. 2b), and the same held at family level (median 0.30 vs 0.26; p = 0.02). Several viral taxa varied consistently between seasons (Fig. 2e): unclassified lineages (log_2_ fold change 0.73), Smacoviridae (2.07), Nanoviridae (1.17), Mimiviridae (1.33) and Crassvirales (0.29) were relatively more abundant in spring, whereas Caudoviricetes (−0.33) and Zobellviridae (−1.61) were enriched in autumn (all BH-adjusted q < 0.05). At the vOTU level, 15 of the 50 vOTUs present in ≥ 2% of samples differed in abundance between seasons; 14 were unclassified vOTUs enriched in spring and one, a Caudoviricetes vOTU predicted to infect *Novosphingobium*, was enriched in autumn (Supp. Fig. S1d). vOTU richness was slightly higher in spring (median 88 vs 79 vOTUs per sample; p = 0.04), whereas Shannon diversity did not differ between seasons or years (Supp. Fig. S1a, b). Viruses of eukaryotes made up a small but seasonally variable fraction of each virome. Their share of per-sample abundance was higher in spring than in autumn (median 1.0% vs 0.4%; mean 5.2% vs 4.2%; Wilcoxon p = 0.008), and spring was higher in each of the three years. The difference came mainly from Mimiviridae, Phycodnaviridae and Nanoviridae, whereas Adintoviridae and Iridoviridae were relatively more abundant in autumn. The repeated occurrence of major viral groups across seasons and years therefore indicates that river biofilms support recurring viral lineages, while strong sample-level turnover and a consistent seasonal shift in their relative abundances indicate that seasonality, rather than interannual variation, is the dominant temporal axis of the biofilm virome.

Taxonomic novelty was substantial across the dataset based on the assembled vOTUs. Although 84.9% of vOTUs were classified at phylum and class level, only 5.2% were assigned to an order and 6.9% to a family (Fig. 2c). This pattern confirms that river biofilms remain a major reservoir of viral dark matter. Placing the classified lineages within a taxonomy-based dendrogram showed that they span twelve viral phyla, with Uroviricota and Nucleocytoviricota accounting for the greatest number of families, while many additional lineages could only be assigned to higher ranks (Fig. 2d). Together, these results show that national river biofilm viromes combine recurring viral groups, high sample-level turnover and extensive taxonomic novelty, highlighting both their ecological structure and poor representation in current viral reference databases.

### Measured water-quality gradients provide limited but biologically interpretable context for virome structure

River biofilm viruses are expected to respond to physicochemical conditions primarily through their microbial hosts, rather than as free-living organisms responding directly to the environment. We therefore tested whether broad patterns in biofilm virome composition were associated with measured water-quality gradients that are known to influence microbial growth, biofilm metabolism and host-community assembly. PCoA based on Bray–Curtis dissimilarity revealed substantial overlap among samples, with spring and autumn samples largely co-occurring in ordination space despite the significant but small seasonal effect (Fig. 3a). Fitted environmental vectors indicated statistically detectable but weak associations with virome structure, with pH showing the strongest association (R^2^ = 0.066), followed by DOC (R^2^ = 0.051), nitrate-N (R^2^ = 0.046), conductivity (R^2^ = 0.044) and dissolved oxygen (R^2^ = 0.021; all BH-adjusted q < 0.05), whereas water temperature, orthophosphate-P and ammoniacal-N were not associated with ordination structure. These results indicate that measured water-quality variables capture only a small proportion of virome variation, consistent with biofilm viral communities being shaped by multiple interacting processes, including host ecology, hydrology, seasonality, spatial history and unmeasured chemical or biological gradients.

**Figure 3.**
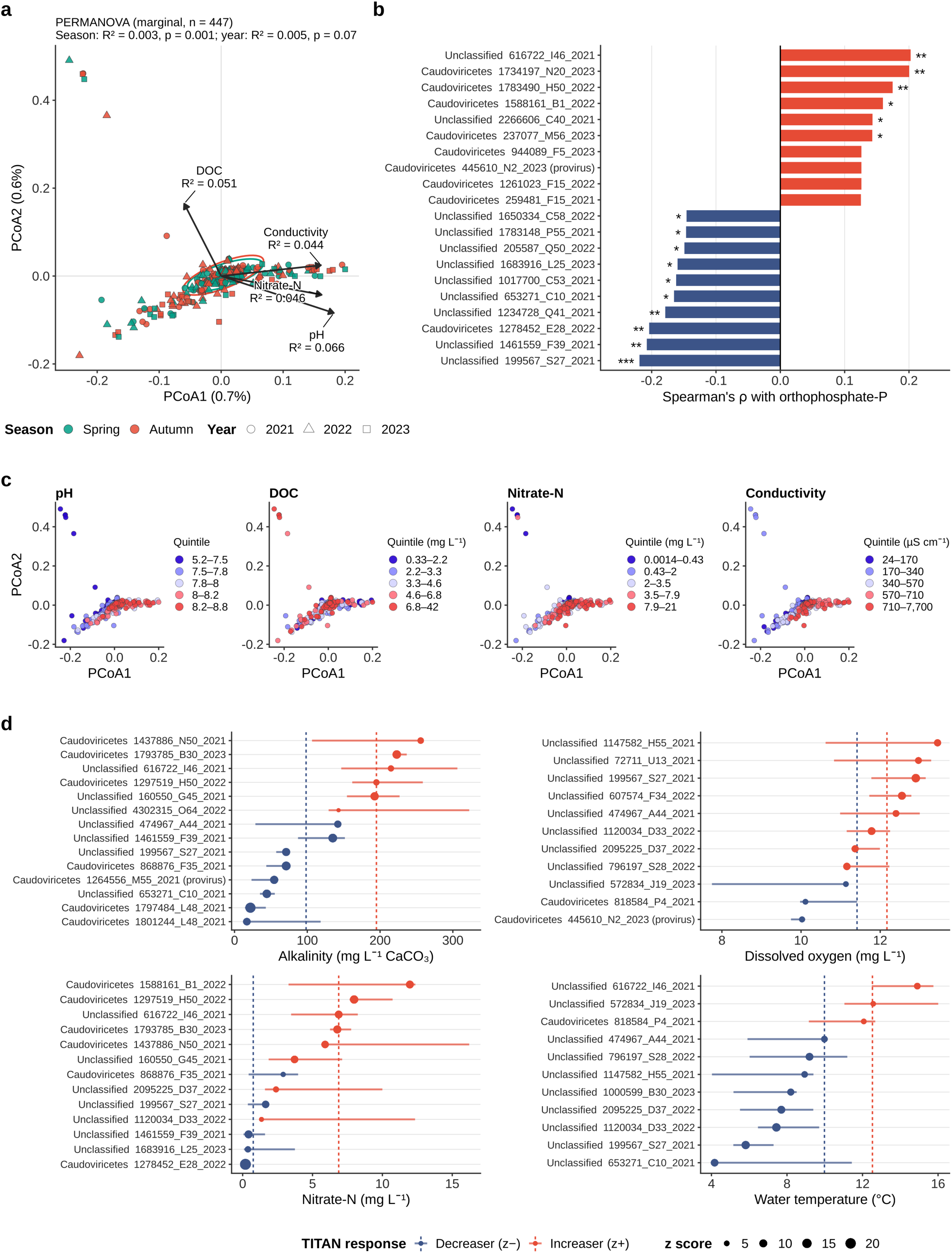
Environmental gradients and seasonality associated with RSN virome structure. (a) PCoA of Bray–Curtis dissimilarities (Hellinger-transformed vOTU abundances; n = 439 samples with complete water chemistry) coloured by season and shaped by year, with 95% ellipses per season and fitted vectors for the four strongest water-quality variables (envfit R^2^). The marginal PERMANOVA for season and year (all 447 samples) is given above the panel. (b) vOTUs with the strongest positive and negative Spearman correlations with orthophosphate-P (BH-adjusted * q < 0.05, ** q < 0.01). (c) The same ordination coloured by sample quintiles of each gradient on a shared low-to-high scale; legends give the value range of each quintile. (d) TITAN change points (median and 5–95% bootstrap interval) for the eight highest-scoring decreasing (z−) and increasing (z+) indicator vOTUs along alkalinity, dissolved oxygen, nitrate-N and water temperature gradients (purity and reliability ≥ 0.95); point size shows the z score and dashed lines the community-level (sum z) change points.

Colouring the ordination by each of the four strongest gradients did not reveal strong separation of samples along any single environmental axis (Fig. 3c); the main cloud of samples spanned the full range of each variable. The distinct group of samples at the margin of ordination space came from a single site sampled in five of the six campaigns, draining a heathland–woodland catchment on siliceous geology. These samples combined low pH, alkalinity, conductivity and nitrate-N with high DOC, and shared many vOTUs across years and seasons despite the extreme sparsity of the dataset, indicating a persistent, site-specific virome. This suggests that the river biofilm virome is unlikely to be explained by any single measured physicochemical variable. Instead, the data support a more complex ecological model in which water quality provides part of the environmental context within which viral communities assemble but does not by itself resolve the drivers of virome composition.

At the vOTU level, some viral populations nevertheless showed weak but detectable associations with nutrient gradients. 23 vOTUs were significantly correlated with orthophosphate-P after BH correction, with Spearman’s ρ reaching approximately ±0.2 (Fig. 3b). Most responsive vOTUs were unclassified or assigned only to Caudoviricetes, indicating that environmentally responsive viral populations include lineages that remain poorly resolved taxonomically. TITAN further identified vOTUs with abrupt change points in abundance and frequency along alkalinity (20 decreasing and 6 increasing), dissolved oxygen (3 and 14), nitrate-N (5 and 9) and water temperature (13 and 3) gradients (Fig. 3d). Community-level change points for decreasing and increasing vOTUs occurred at approximately 98 and 195 mg L^−1^ CaCO_3_ for alkalinity, 11.4 and 12.2 mg L^−1^ for dissolved oxygen, 0.8 and 6.9 mg L^−1^ for nitrate-N, and 10.0 and 12.5 °C for water temperature. Because only 50 vOTUs were sufficiently prevalent for TITAN and the same vOTUs responded to several correlated gradients, these population-level responses should be interpreted cautiously. Of the 37 indicator vOTUs, 14 also differed in abundance between seasons, and indicators occurred at a median of 10 sites (range 5–25), so some change points are likely to reflect seasonal turnover or the chemistry of a few sites. However, they suggest that although bulk community ordination is only weakly structured by measured water chemistry, individual viral populations may still respond to specific environmental gradients.

### Host-virus associations reveal modular ecological structure linked to water quality

Because viral composition is expected to reflect host ecology and did not reveal strong associations with water-quality gradients, we next integrated predicted host associations with viral taxonomy and environmental gradients. Abundance-weighted host–virus lineage summaries showed that the RSN virome was linked primarily to metagenome-assembled genomes (MAGs) from Pseudomonodota, Cyanobacteriota and Bacteroidota, with additional interactions with Bacillota, Actinomycetota and other phyla (Fig. 4a). Most abundant viral associations were assigned to Caudoviricetes or remained unclassified at family level, again emphasising that ecological interpretations potentially require the combination of taxonomic, host-linked and functional information rather than relying on viral taxonomy alone. Host prediction was limited to bacteria and archaea, because iPHoP, which integrates homology-, CRISPR-spacer- and composition-based predictors (see Methods), does not predict eukaryotic hosts. Viruses of eukaryotes (1,716 vOTUs, mainly Adintoviridae, Mimiviridae, Phycodnaviridae and Iridoviridae) accounted for about 5% of total virome abundance and were therefore not assigned hosts.

**Figure 4.**
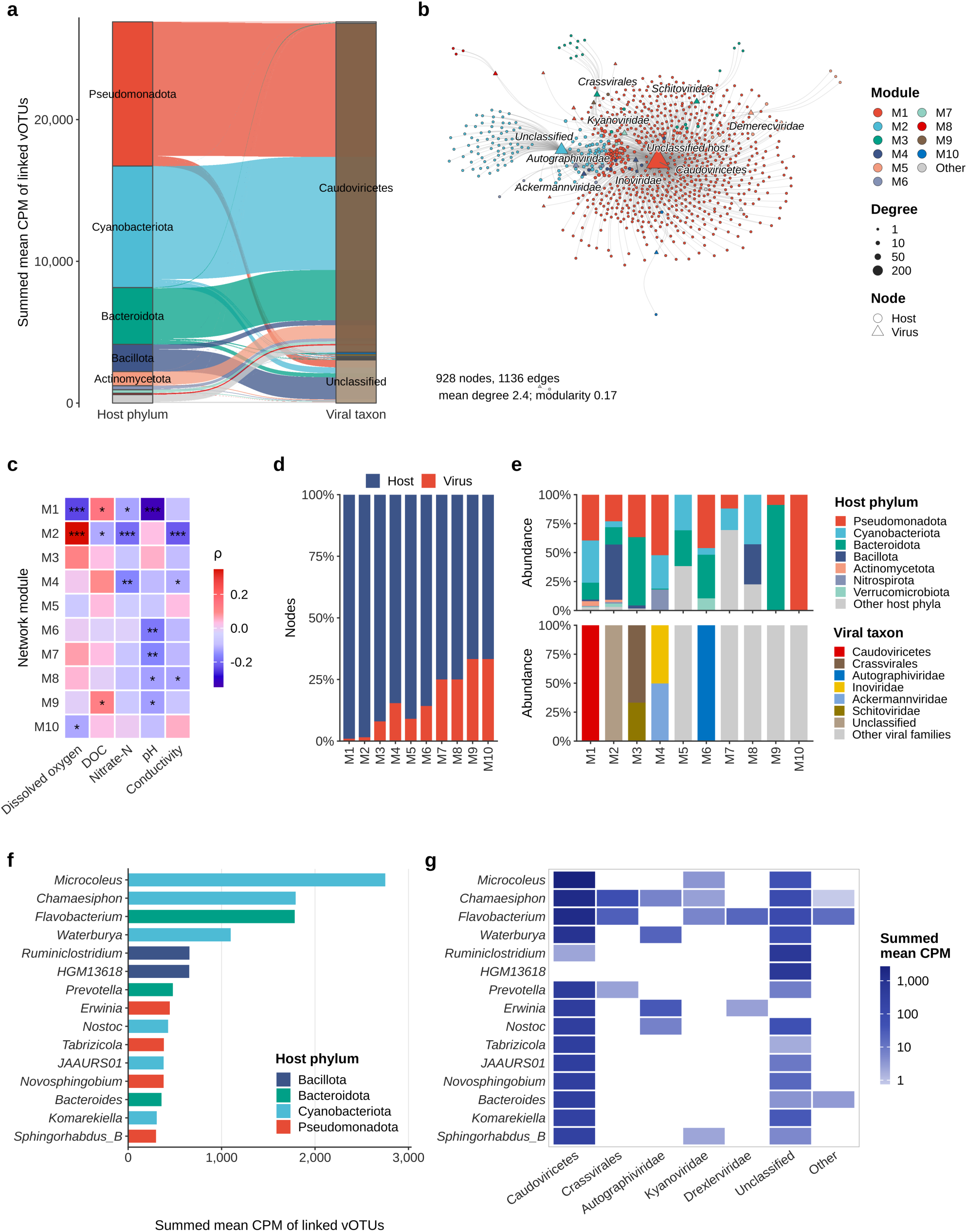
Host–virus associations and environmental structure of the RSN virome. (a) Alluvial summary of predicted host phyla and viral taxa, weighted by summed mean CPM of linked vOTUs (iPHoP confidence ≥ 90). Viruses of eukaryotes are excluded from all panels. (b) Bipartite host genus–viral taxon network (Fruchterman–Reingold layout) coloured by Louvain module (ten largest shown); node size is proportional to degree and hub taxa are labelled. (c) Spearman correlations between module viral abundance and water-quality variables across samples (BH-adjusted * q < 0.05, ** q < 0.01, *** q < 0.001). (d) Proportion of host and viral nodes in each module. (e) Abundance-weighted host-phylum (top) and viral-taxon (bottom) composition of each module. (f) Dominant predicted host genera ranked by the summed mean CPM of linked vOTUs. (g) Viral taxa linked to the dominant host genera, on a log_10_ colour scale.

The overall predicted host–virus association network contained 928 nodes and 1136 edges, with an average degree of 2.4, low density (0.003) and 12 Louvain modules (Fig. 4b). The global modularity value was 0.17, indicating weak, yet non-random compartmentalisation in a large and sparse network, with a few viral taxa (notably Caudoviricetes and unclassified viruses) acting as hubs connecting many host genera. Module-level viral abundances were significantly correlated with water-quality variables after BH correction (Fig. 4c). The largest, Caudoviricetes-dominated module (M1) was negatively correlated with pH (ρ = −0.33) and dissolved oxygen (ρ = −0.24) and positively with DOC (ρ = 0.14), whereas the module dominated by unclassified viruses (M2) showed the opposite response to dissolved oxygen (ρ = 0.35) and was negatively correlated with conductivity and nitrate-N. Because M1 and M2 are centred on the Caudoviricetes and unclassified-virus nodes, their opposite associations with dissolved oxygen largely mirror the seasonal shift between these groups (Fig. 2e). These contrasting responses support the idea that environmental filtering acts through distinct host–virus compartments rather than uniformly shifting the entire virome.

All major modules contained both host and viral nodes, although host genera dominated numerically (Fig. 4d). Host composition differed among compartments, with modules enriched for Pseudomonodota, Cyanobacteriota, Bacteroidota or Bacillota, while viral compartments were dominated by Caudoviricetes, unclassified viruses, Inoviridae and Ackermannviridae, Crassvirales and Schitoviridae, and Autographiviridae (Fig. 4e). Dominant predicted host genera included *Microcoleus*, *Flavobacterium*, *Chamaesiphon*, *Waterburya*, *Ruminiclostridium*, *Prevotella*, *Erwinia*, *Nostoc*, *Tabrizicola* and *Novosphingobium* (Fig. 4f). *Flavobacterium* and *Chamaesiphon* were linked to the largest numbers of vOTUs (457 and 268, respectively), whereas *Microcoleus* ranked first by abundance with 114 linked vOTUs, and the high ranks of *Ruminiclostridium*, HGM13618 and *Thomasclavelia* each reflected only one or two abundant vOTUs.

Viral family associations differed among these host genera (Fig. 4g), with some hosts, such as *Flavobacterium* and *Chamaesiphon*, linked to a broad set of viral families and others, such as *Ruminiclostridium* and *Thomasclavelia*, dominated by a single viral association. Collectively, these findings suggest that host associations may link water quality and virome structure, a hypothesis that host-resolved analyses could test directly.

### Viral life-history traits and AMGs indicate functional coupling to biofilm metabolism

We evaluated whether the ecological structure of the RSN virome was reflected in viral traits and metabolic potential. Genome size was strongly associated with predicted gene content (R^2^ = 0.88, ≈ 1.3 genes kb^−1^; Fig. 5a), consistent with internally coherent viral genome predictions. Only 458 vOTUs (1.2%) were identified as proviruses and classified as temperate; these had larger genomes than lytic vOTUs (median 18.0 vs 7.6 kb; Wilcoxon p < 0.001; Fig. 5b), consistent with many lytic vOTUs representing genome fragments. Provirus-derived vOTUs made up a small share of abundance across all major viral taxa (Fig. 5c). Viruses of eukaryotes were excluded from lifestyle comparisons (Fig. 5c and 5d).

**Figure 5.**
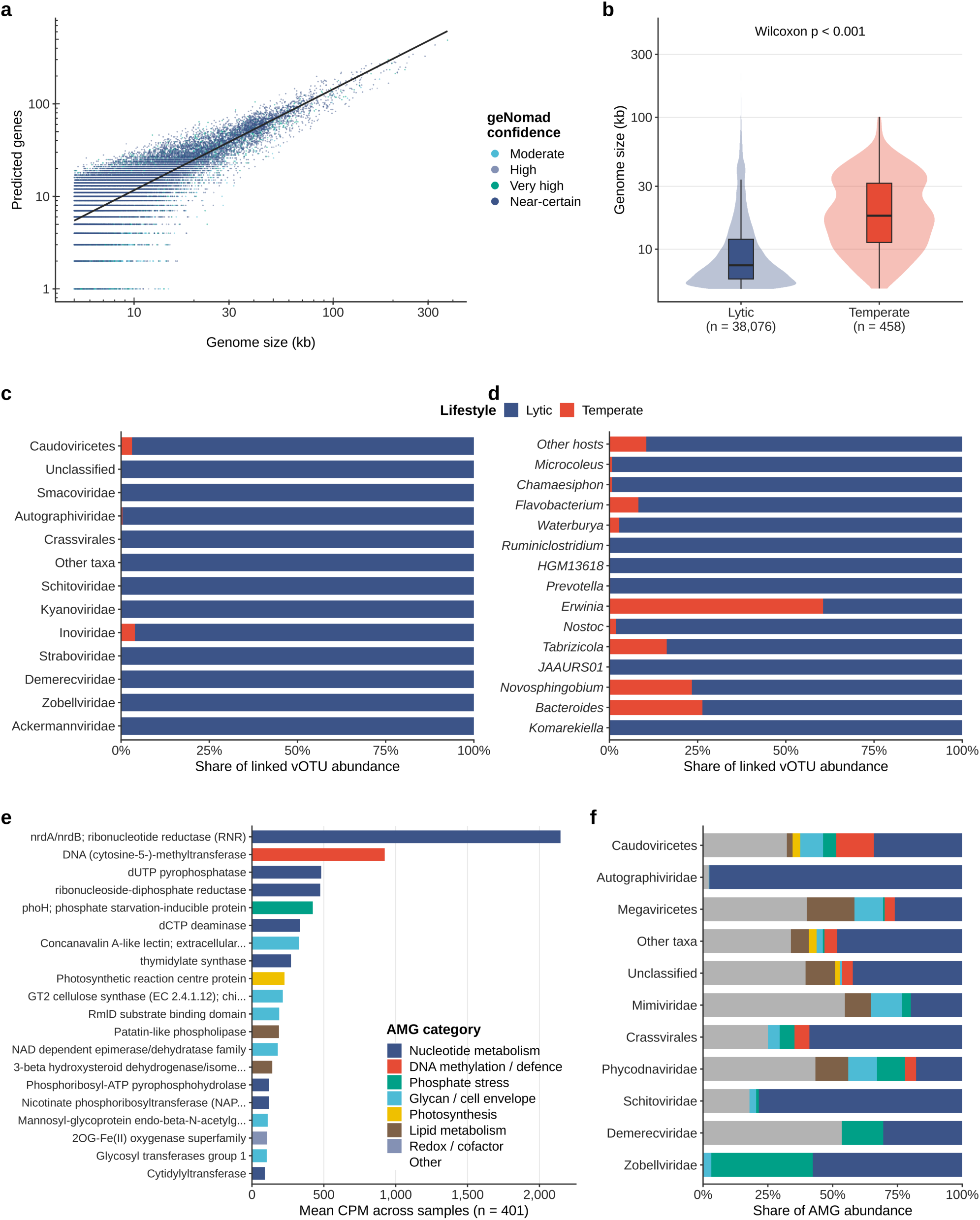
Viral genome traits, lifestyle predictions and auxiliary metabolic genes in RSN viromes. (a) Relationship between genome size and predicted gene content for all 38,534 vOTUs, coloured by geNomad identification confidence (log–log axes; line, ordinary least squares fit). (b) Genome-size distributions of lytic and temperate (provirus) vOTUs. (c) Share of abundance attributable to lytic and temperate vOTUs within the dominant taxa of prokaryotic viruses. (d) As (c) for the dominant predicted host genera. (e) Top 20 AMG functions ranked by mean CPM across 401 samples (samples with DRAM-v AMG output), coloured by functional category. (f) Functional-category composition of AMGs within the dominant encoding viral taxa.

Lifestyle composition varied more strongly across predicted host genera than across viral taxonomic categories (Fig. 5d). Most dominant host-linked viral assemblages were lytic-dominated, but selected genera, including *Erwinia* (61% temperate), *Bacteroides* (26%), *Novosphingobium* (23%) and *Tabrizicola* (16%), showed larger temperate fractions. This pattern is consistent with host ecology shaping viral life-history strategies. Rather than suggesting that lysogeny may be a direct response to any single anthropogenic pressure, these results indicate that viral lifestyle variation is shaped by host context and should be considered an emergent trait of the wider biofilm community.

To understand the functional roles of the virome, we characterised the functional potential of the RSN virome. Functional annotation identified a diverse set of AMGs across the virome (Fig. 5e). The dominant AMG signal was *nrdA*/*nrdB* ribonucleotide reductase, followed by DNA cytosine-5-methyltransferase, dUTP pyrophosphatase, ribonucleoside-diphosphate reductase, *phoH*, dCTP deaminase, a concanavalin A-like lectin, thymidylate synthase, GT2 cellulose synthase-related functions, RmlD and NAD-dependent epimerase/dehydratase domains, photosynthetic reaction centre protein, patatin-like phospholipase and other functions. Nucleotide-metabolism genes accounted for most of the dominant AMG abundance. These AMGs map onto processes that are highly relevant to biofilm ecology such as nucleotide biosynthesis for viral replication, methylation and defence-related interactions, phosphate stress, cell-envelope and glycan modification, lipid metabolism, redox processes and photosynthetic energy capture. Simultaneously, it must be noted that nrdA/nrdB, dUTPase, dCTP deaminase, thymidylate synthase and methyltransferases are generally considered replication- or defence-associated rather than metabolic AMGs.^27^

Furthermore, AMG composition differed among encoding viral taxa (Fig. 5f). Autographiviridae and Schitoviridae AMG repertoires were dominated by nucleotide-metabolism genes, Zobellviridae and Demerecviridae carried a larger share of phosphate-stress (*phoH*) genes, and Mimiviridae, Megaviricetes and Phycodnaviridae carried relatively more glycan/cell-envelope and lipid-metabolism genes, which are likely part of the core gene content of these giant viruses rather than auxiliary genes. These also indicate that functional potential is not evenly distributed across the virome but is structured by viral lineage and, by implication, host association. Importantly, the abundance of photosynthesis-related and glycan-modification AMGs observed in our dataset is particularly relevant for benthic biofilms, where phototrophs, heterotrophs and extracellular polymeric substances form a tightly coupled metabolic matrix. Collectively, these results support the hypothesis that river biofilm viruses are not simply passive markers of microbial community composition. They highlight the notion that the virome encodes metabolic functions that may influence host physiology and biofilm biogeochemical processes.

## Discussion

River biofilms are foundational components of freshwater ecosystems, forming spatially structured, matrix-enclosed communities of bacteria, algae, fungi and other microorganisms. These microbial assemblages regulate nutrient cycling, organic matter processing, primary production and water-quality transformations, yet the viral communities embedded within them remain poorly characterised, particularly at national scales. Previous work on stream biofilms has shown that viral diversity can be strongly linked to bacterial community composition and structured across seasons and stream gradients,^28^ supporting the view that biofilm viruses are ecologically organised rather than randomly assembled. Our study extends this perspective to a national river surveillance network, showing that England’s river biofilm viromes are not simply transient collections of viral particles moving downstream, but highly novel and seasonally structured communities with host-linked and functional organisation that reflects the state of the biofilm environment.

A central outcome of our study is the establishment of a national-scale baseline for river biofilm viral diversity. Across the RSN metagenomes, we recovered more than 38,000 vOTUs, the vast majority of which could not be resolved to low taxonomic ranks. This degree of novelty is consistent with broader freshwater virome studies,^29^ which show that freshwater viral diversity remains substantially under-characterised and poorly represented in reference databases. Global freshwater virome analyses have similarly highlighted extensive unclassified viral diversity, reinforcing the need for ecosystem-specific viral catalogues.^30^ Our finding that Caudoviricetes and unclassified viral lineages repeatedly dominate across years and seasons therefore fits with the wider freshwater virome literature, while showing that this pattern also holds within benthic river biofilms at national scale.

This taxonomic uncertainty does not make the biofilm virome ecologically uninterpretable. The repeated recovery of major viral groups across years and seasons, combined with pronounced sample-level turnover, indicates that river biofilms contain both recurring viral groups and substantial local heterogeneity.^31^ Within this heterogeneity, seasonality emerged as the dominant temporal signal: season, but not year, explained a significant share of taxon-level composition, and within-site turnover between spring and autumn exceeded turnover between years. Spring biofilms were relatively enriched in unclassified lineages and in small ssDNA viruses (Smacoviridae, Nanoviridae) and giant viruses of eukaryotes (Mimiviridae), whereas autumn biofilms were enriched in Caudoviricetes, the tailed phages of bacteria. This shift coincided with warmer, less oxygenated and more phosphate-rich autumn conditions and plausibly tracks seasonal succession of biofilm hosts, from spring communities rich in diatoms and other eukaryotic algae towards more heterotrophic, bacteria-dominated autumn biofilms. Consistent with this, viruses of eukaryotes made up a larger share of the virome in spring than in autumn, suggesting that part of the seasonal signal is tied to eukaryotic hosts that our genome-based host prediction could not capture. This pattern is biologically plausible because biofilms are spatially structured habitats in which host populations, extracellular polymeric substances, redox gradients, nutrient fluxes and hydrological exposure vary across fine scales. Experimental work has shown that phages can become trapped within biofilm matrices and remain infectious, influencing colonisation dynamics and microbial organisation.^12^ The river biofilm virome therefore may be better understood as an integrated record of host–virus interactions, a hypothesis that time-resolved sampling could test in the future.

Anthropogenic pressures influence river ecosystems through changes in nutrient loading, organic matter availability, oxygen dynamics, chemical exposure, hydrology and microbial community organisation. For biofilm viruses, these effects are likely to be expressed through host availability, host physiology and infection context, rather than through direct exposure to physicochemical conditions alone. The measured water-quality gradients in our study therefore provide a biologically meaningful framework for linking river environmental condition to viral community structure, host associations and functional potential. However, whole-community ordination showed that pH, dissolved organic carbon, conductivity, nitrate-N and dissolved oxygen each explained at most 7% of virome variation, with the strongest pH and DOC contrasts coming from a single distinctive heathland site, and no single measured gradient produced strong separation of samples.

The limited explanatory power we observed is an important ecological result. It suggests that river biofilm viromes are not governed by a simple one-variable environmental axis. Instead, their structure likely emerges from the combined effects of host-community composition, hydrological connectivity, seasonal dynamics, local biofilm history, unmeasured chemical mixtures and stochastic colonisation processes. The extreme sparsity of the vOTU table, in which no vOTU occurred in more than 8.3% of samples, also means that vOTU-level dissimilarities are close to saturation, which limits the variance any predictor can explain; aggregating to viral taxa recovered a clearer seasonal signal. Such complexity is expected in national-scale river networks, where environmental gradients overlap and where biofilms integrate biological and physicochemical signals over time. Measured water quality therefore explains a small fraction of vOTU-level variation, consistent with multiple interacting filters, only some of which are captured by routine water-quality variables.

At the level of individual viral populations, some vOTUs nevertheless showed environmentally responsive behaviour. Orthophosphate-P was associated with both positively and negatively responding vOTUs, and TITAN identified candidate indicator viral populations with abrupt change points along alkalinity, dissolved oxygen, nitrate-N and temperature gradients. These responses should be interpreted cautiously because effect sizes were small, only the most prevalent vOTUs could be tested, and many responsive vOTUs remain taxonomically unresolved. Also, several indicator vOTUs also differed between seasons. However, they suggest that even when bulk virome ordination is weakly structured, individual viral populations may still track specific physicochemical conditions. This has direct relevance for biomonitoring because the strongest viral signal may lie not in broad community separation, but in the behaviour of particular indicator populations, host-linked modules or functional traits.

The host-association findings suggest a route by which water quality could shape virome structure. Predicted hosts were dominated by plausible river biofilm lineages, including Cyanobacteriota, Bacteroidota and Pseudomonodota, with prominent host-linked genera such as *Microcoleus*, *Chamaesiphon* and *Flavobacterium*. These groups occupy distinct ecological roles in biofilms, including phototrophic primary production, organic matter degradation, opportunistic growth and nutrient transformation. The host-linked structure of the RSN virome is consistent with stream-biofilm evidence showing that viral diversity is connected to bacterial community composition.^28^ In our dataset, predicted virus–host associations formed a sparse network with weak modularity, organised around a few highly connected viral lineages. Module-level viral abundances correlated with several water-quality variables. However, the largest modules were centred on Caudoviricetes and unclassified viruses, so these correlations largely mirror the seasonal shift between the two groups. Whether environmental filtering acts through distinct host–virus compartments, rather than uniformly across the virome, therefore, remains a hypothesis that host-resolved analyses at the vOTU level could test.

The modular structure of the host–virus network provides a biologically meaningful way to interpret virome complexity.^32^ Different modules contained distinct combinations of host phyla and viral groups, including Pseudomonodota-, Cyanobacteriota- and Bacteroidota-associated compartments linked to Caudoviricetes, Inoviridae, Ackermannviridae, Crassvirales, Schitoviridae, Autographiviridae and unclassified viruses. Such compartmentalisation may reflect host specificity, microhabitat structure within the biofilm matrix, differences in host physiology or variation in infection strategy.^32^ Rather than indicating a uniform viral response to river condition, these modules suggest that biofilm viromes are organised into ecological sub-networks that may respond differently to changing physicochemical conditions; for example, the Caudoviricetes-dominated and unclassified-virus-dominated modules showed opposite associations with dissolved oxygen. This is especially relevant for river health, because disruption of particular host–virus compartments could alter microbial turnover, phototrophic activity, nutrient processing or organic matter flow.^29^

Viral lifestyle patterns further support a host-context-dependent view of river biofilm viral ecology. Host growth state, density, immunity, physiological stress and local resource availability are all likely to influence whether lytic or temperate strategies are favoured. In our dataset, lytic viruses dominated most viral taxa and host-linked assemblages, although fragmented genomes and our provirus-based definition mean that temperate viruses are likely under-detected. Temperate fractions were elevated in selected host genera, suggesting that lifestyle strategies are embedded within specific host–virus interactions rather than uniformly tracking a single environmental gradient. This interpretation fits the broader biology of biofilms, where matrix structure can alter encounter rates, viral diffusion, host defence and viral persistence.^12^

Our AMG results indicate that river biofilm viruses encode functional potential relevant to microbial metabolism and river biogeochemistry. The strongest AMG signal was *nrdA*/*nrdB* ribonucleotide reductase, alongside other nucleotide-metabolism genes such as ribonucleoside-diphosphate reductase, dUTP pyrophosphatase, dCTP deaminase and thymidylate synthase. Ribonucleotide reductases are widely distributed in viral genomes and have been proposed as useful markers of viral diversity because they support deoxyribonucleotide production for viral genome replication.^33^ Their dominance in the RSN virome is therefore biologically plausible and suggests strong selection for viral control of nucleotide precursor supply during infection. Other AMGs point towards host–virus defence interactions and nutrient-related metabolic modulation. DNA cytosine methyltransferases suggest interactions with host restriction-modification and defence systems,^34^ while *phoH* is commonly interpreted as a viral AMG associated with phosphorus stress or phosphate-related metabolism. These genes indicate functional potential through which viruses may influence the physiological state of their hosts during infection and thereby contribute to biofilm metabolic dynamics.

Several AMGs were particularly relevant to the biofilm mode of life. Glycosyltransferases, lectins, GT2 cellulose synthase-related functions, patatin-like phospholipases and NAD-dependent epimerase/dehydratase functions are associated with cell-envelope modification, extracellular polymer production, lipid metabolism or matrix-associated processes. These functions are important because the extracellular matrix controls biofilm adhesion, protection, nutrient retention, diffusion and spatial organisation. Given that matrix-associated phages can remain infectious and influence microbial colonisation,^12^ the presence of matrix-relevant AMGs in the RSN virome raises the possibility that viruses influence not only host mortality, but also the physical and metabolic architecture of biofilms.

Photosynthesis-related AMGs provide another important biological signal. Cyanophages are well known to carry genes involved in photosynthesis and host metabolic reprogramming, allowing viruses to maintain energy flow during infection.^35^ Our detection of photosynthetic reaction centre proteins, combined with predicted associations to phototrophic hosts such as Cyanobacteriota and genera including *Microcoleus* and *Chamaesiphon*, suggests that similar viral strategies may operate within benthic river biofilms. This is particularly relevant because phototrophs and heterotrophs are tightly coupled in biofilms, and viral modulation of phototrophic metabolism could influence local energy flow, oxygen dynamics and organic carbon supply.

The distribution of AMGs across viral taxa further reinforces the idea that viral functional potential is ecologically structured. Caudoviricetes, Autographiviridae, Mimiviridae, Phycodnaviridae, Schitoviridae, Crassvirales and unclassified viruses carried different combinations of dominant AMG functions. Similar freshwater virome studies have reported viral genes linked to metabolic processes including carbohydrate metabolism, nutrient cycling and host metabolic modulation, indicating that viral functional potential is a recurrent feature of freshwater ecosystems.^29,30^ In the RSN dataset, these functions are embedded within a biofilm-specific ecological architecture, where taxonomic novelty, host associations and matrix-associated metabolism intersect. Because many abundant and environmentally responsive viruses remain taxonomically unresolved, functional and host-linked analyses become essential for biological interpretation.

Collectively, these findings have important implications for river monitoring. Current freshwater assessment frameworks are built primarily around physicochemistry, macroinvertebrates, algal indices, faecal indicators, bacterial communities or targeted chemical surveillance. Viromics adds a complementary layer because viruses are tightly connected to microbial mortality, host physiology, horizontal gene transfer and metabolic reprogramming. However, our results also caution against treating the virome as a simple diagnostic readout. The most defensible interpretation is that biofilm viromes integrate multiple ecological signals, including season, measured water quality, host-community structure, viral lifestyle and functional gene content. The strong seasonal signal also implies that virome-based monitoring should compare samples within, rather than across, seasons. For monitoring applications, viromics is therefore likely to be most powerful when combined with water chemistry, microbial host profiles and catchment information, rather than used in isolation.

Several key limitations remain in leveraging viromes for freshwater assessment. Host associations were computationally predicted and should be interpreted as hypotheses of infection rather than confirmed virus–host pairs. Viral taxonomy remains incomplete, limiting biological interpretation for many vOTUs, including some environmentally responsive populations. The extreme sparsity of vOTU occurrences restricted indicator analyses to a small number of prevalent vOTUs. AMG annotation identifies functional potential but does not demonstrate expression or activity. Future studies will need additional approaches such as metatranscriptomics, proteomics, viral tagging, proximity-ligation (Hi-C) or single-cell approaches, long-read assembly of complete CRISPR arrays and proviruses or targeted validation to confirm active viral metabolic reprogramming and infection links. Finally, river environmental gradients are correlated with each other and with season, making it difficult to assign causality to individual variables. These limitations are inherent to national-scale field viromics, but they also define clear priorities for future work, particularly linking viromes to matched host-community profiles, chemical mixtures, hydrological data and time-resolved measurements of viral activity.

Our national-scale findings reposition river biofilm viromes as a critical but overlooked layer of freshwater biodiversity. They are taxonomically novel, spatially heterogeneous and seasonally structured, organised around plausible microbial hosts and encoding functions relevant to nucleotide metabolism, host defence, nutrient stress, photosynthesis and biofilm matrix processes. The strongest ecological interpretation is not that water quality alone explains virome structure, but that viral communities encode the combined imprint of season, physicochemical context, host-community organisation and metabolic interaction within the biofilm matrix. Anthropogenic pressures are likely to act through this ecological context, which future work linking viromes to pressure data could test. Incorporating viromics into river biofilm ecology therefore offers a new route to understanding freshwater health, microbial function and ecosystem response under increasing environmental change.

## Data Availability

Raw sequencing reads have been deposited in the European Nucleotide Archive (ENA) at EMBL-EBI under accession number PRJEB85861. The DiscoVir pipeline code and downstream analysis scripts used in this study are available at https://doi.org/10.5281/zenodo.23601835 and https://zenodo.org/records/23602010. Processed data tables, including vOTU abundance matrices, host-prediction summaries and AMG annotations, are provided in the Supplementary table 1.

## Code Availability

Downstream R scripts used to generate the figures, seasonal statistics and TITAN analyses (00_prep.R, 01_stats.R and fig1–5, figS1 scripts) are available at https://zenodo.org/records/23602010.

## Author contributions

S.B.B.: conceptualisation, methodology, formal analysis, visualisation, validation, investigation, data curation, writing-original draft, project administration, and supervision. A.C.T.: conceptualisation, methodology, resources, validation, project administration, and writing-original draft. J.W.: funding acquisition, resources and writing-review. K.W.: funding acquisition, resources, validation, writing-review, and supervision. D.S.R.: conceptualisation, project administration, funding acquisition, resources, validation, writing-original draft, and supervision.

## Declaration of competing interest

The views expressed in this paper are those of the authors and do not necessarily reflect the views of the Environment Agency. The authors declare that they have no known competing financial interests or personal relationships that could have appeared to influence the work reported in this paper.

## Acknowledgements

We thank Laura Hunt for collating the RSN metadata. We also thank Dr Jonathan Porter, Sean Butler and Alan Wan from the Environment Agency’s National Monitoring Laboratories for their support in collating and shipping the biofilm samples for further processing.

## Funding

This work was funded by the Environment Agency under research project SC220034. ACT, SBB and DSR were supported by Natural Environment Research Council (NERC) grant NE/Z000173/1. KW and JW were supported by NERC grant NE/X015777/1.

## Supplementary figure legend

Supplementary Figure S1. Diversity, sparsity and seasonally responsive vOTUs. (a) vOTU richness and (b) Shannon diversity per sample by year and season (Wilcoxon test for season, Kruskal–Wallis test for year). (c) Distribution of vOTU occurrence across the 447 samples (log–log); dashed lines mark 2% and 10% prevalence. (d) Seasonal differential abundance of the 50 vOTUs detected in ≥ 2% of samples (Wilcoxon, BH-adjusted); labelled vOTUs have q < 0.05.

## Notes

https://doi.org/10.5281/zenodo.23601835

https://zenodo.org/records/23602010

### Competing Interest Statement

The authors have declared no competing interest.

